# Dengue virus capsid protein interaction with nucleic acids

**DOI:** 10.1101/2025.07.14.664594

**Authors:** Nelly M. Silva, Ana S. Martins, Nina E. Karguth, Maria J. Sarmento, Francisco J. Enguita, Roland G. Huber, Ivo C. Martins, Nuno C. Santos

**Affiliations:** GIMM – Gulbenkian Institute for Molecular Medicine, Av. Prof. Egas Moniz, 1649-035 Lisbon, Portugal; Faculdade de Medicina, Universidade de Lisboa, Av. Prof. Egas Moniz, 1649-028 Lisbon, Portugal; Bioinformatics Institute (BII), Agency for Science, Technology and Research (A*STAR), Singapore 138671, Singapore

**Author notes:** **Correspondence:** <u>Ivo C. Martins</u>, GIMM – Gulbenkian Institute for Molecular Medicine/ Faculdade de Medicina, Universidade de Lisboa, Av. Prof. Egas Moniz, 1649-028 Lisbon, Portugal.; <u>Nuno C. Santos</u>, GIMM – Gulbenkian Institute for Molecular Medicine/ Faculdade de Medicina, Universidade de Lisboa, Av. Prof. Egas Moniz, 1649-028 Lisbon, Portugal. **Declaration of Competing Interest** The authors declare no conflict of interest.

**Keywords:** *Flavivirus*, nucleic acids, capsid protein, protein-DNA interaction, liquid-liquid phase separation

## Abstract

Dengue virus (DENV) transmission has greatly increased in the last decade, in part due to the geographical expansion of its vector, the *Aedes* spp. mosquitoes. These arthropods are now found even in temperate climates, including Europe, where outbreaks have already occurred. A better understanding of the virus life cycle is essential, as it may enable the development of specific treatments or therapies which are currently lacking. However, recent breakthroughs concerning the biologically relevant viral capsid (C) protein structure and function are encouraging. It is now clear the C protein binds both to host lipid droplets and to the viral genome, a set of interactions that is crucial for viral encapsidation and replication. Previously, we studied DENV C protein interaction with lipid droplets, at the molecular level. Here, we examine how the association of DENV C protein with the viral genome occurs. Using DENV C protein and single-stranded DNA (ssDNA) sequences analogous to previously identified important regions of the viral genome, we biophysically characterize their interactions. A decrease in fluorescence intensity and lifetime, an increase in both anisotropy and hydrodynamic diameter, as well as changes in the protein’s secondary structure, were observed upon binding to ssDNA. These changes are consistent with molecular condensation and raise the possibility of liquid-liquid phase separation involvement in DENV C-nucleic acid complex formation. These findings may be useful for the development of future therapeutic strategies based on nucleic acids, not only for DENV but also for other related viruses.

## 1. Introduction

Dengue virus (DENV) is a member of the *Flaviviridae* family, which comprises important human pathogens, such as Zika virus (ZIKV), West Nile virus (WNV), yellow fever virus (YFV), Japanese encephalitis virus, and tick-borne encephalitis virus. Although various modes of transmission have been identified, DENV and ZIKV are mainly transmitted via *Aedes* spp. mosquito bites. The geographical spread of these viral vectors has dramatically increased over the last few years [1]. Due to climate change, several regions of the world (including Europe and North America) are becoming suitable for the establishment of these flavivirus’ vectors [2,3]. Combined with the globalization of trade and travel (facilitating the spread of both the vector and the virus itself), it is expected that this trend towards an even larger global expansion will continue [4].

DENV virion consists of a spherical particle formed by a lipid bilayer associated with two structural proteins, the membrane (M) and envelope (E) proteins. They surround the viral nucleocapsid, which is composed by the viral genome, a positive-sense single-stranded RNA (ssRNA), and by the structural capsid (C) protein [5]. *Flavivirus* cryogenic electron microscopy structures show poor nucleocapsid density [5–7], which might be explained by different scenarios: *i*) the core has an asymmetric structure, lacking organization; *ii*) the core does not have a single structure; or, *iii*) the icosahedral orientation is not synchronized with the external structure [5,8–12]. Moreover, the amorphous nucleocapsid may also be justified by the tendency of DENV C protein to bind the RNA genome indiscriminately [5,10,11,13]. Regardless of how it is exactly organized, there is a body of evidence showing that DENV C interacts directly with the viral RNA, not only in the mature virion structure, but also during the viral assembly and encapsidation processes [5,14].

DENV C is composed of 100 amino acid residues, and forms homodimers stabilized by electrostatics and hydrophobic interactions [15–17]. Monomers are characterized by four α-helices (α1 to α4) connected by short loops, and a transiently disordered N-terminal region [8,15]. Based on the X-ray crystallography and nuclear magnetic resonance structures [15,18,19], it was proposed that the observed asymmetric charge distribution is intimately associated with the C protein function: the α4-α4’ region is likely to interact with the viral genome, while the α2-α2’ was shown to interact with host lipid systems [8,13,15,17,20–24]. It is also possible that the positively charged amino acid residues of the N-terminal region play a role in the binding to RNA [25], since the N-terminal region of members of the *Hepacivirus* genus is already considered a flexible RNA-binding domain [23,25] and, in parallel, given the RNA negative charge, electrostatics charge considerations would favor such an interaction (even if not of a specific nature). Recent findings suggest that the highly electropositive surface across the entire protein indicates a complex assembly mechanism, not solely dependent on charge asymmetry [26,27].

As previously mentioned, the C protein is extremely important for viral assembly by ensuring viral genome encapsidation [5,14,28]. It is responsible for the recruitment of newly synthesized viral genome to form the nucleocapsid, which subsequently buds from the endoplasmic reticulum (ER), acquiring the lipid bilayer together with the remaining viral structural proteins (E and prM) [29,30]. Mutations deleting part of the α3 helix impaired propagation and infectivity [28]. Furthermore, single-point mutations on residues located in the α2-α2’ (L50S and L54S) and α4-α4′ (L81N and I88N) regions affected the structural stability of the protein and the RNA-capsid protein interaction [31]. More recently, it was confirmed that the neutralization required to generate the nucleocapsid is likely to be restricted to α4-α4′ positive residues, R85 and K86, rather than being unspecific [32]. Indeed, mutations of those residues do not affect the protein functionality but increase the formation of defective viral particles [32,33].

During encapsidation, the C protein also promotes RNA folding to form the nucleocapsid [29] and is involved in genome cyclization, which is crucial for viral replication [34]. After viral translation, the C protein transiently interacts with the membrane and may initiate viral assembly by interacting with viral RNA, as well as other C proteins [25,34,35]. Even though the phylogenetic proximity with alphavirus, flavivirus nucleocapsids are poorly ordered, with a remarkedly different assembly [13,36–39]. While *Flavivirus* virions are assembled close to the ER, alphaviruses assemble on the plasma membrane [40]. Even comparing with viruses that share an assembly at the ER level or close to it, there are differences, demonstrating that the mechanisms can be complex and diverse [40].

Overall, due to its complexity, the interaction of C protein and nucleic acids is less understood. Nevertheless, previous studies have identified DENV C as a supercharged protein with potential as a nucleic acid delivery system. In fact, DENV C effectively facilitated the delivery of single-stranded DNA (ssDNA) across various mammalian cell models, suggesting genome translocation as an additional role for DENV C [41]. Peptides derived from DENV C were also evaluated and found capable of entering cells [41,42], presenting high affinity for nucleic acids, as determined by tryptophan residue fluorescence quenching [42]. The assembly of nucleocapsid-like particles (NCLP) in solution has also been studied *in vitro*, revealing oligonucleotide length dependence, where a 5-mer oligonucleotide was found to be sufficient to induce assembly. Due to the small size of this oligonucleotide, its secondary structure seems to have little influence on NCLP formation [26].

Herein, we report our findings regarding the interaction of DENV C with nucleic acids derived from the viral genome. We used ssDNA sequences analogous to important regions of the genome [43], and employed spectroscopic techniques to study the changes from a protein point of view. The results obtained – including changes in fluorescence properties, protein secondary structure and hydrodynamic diameter – may be consistent with a mechanism involving liquid-liquid phase separation, a process recently suggested for DENV C in the presence of nucleic acids, under different experimental conditions [30]. These findings provide insights not only into DENV C but may also have a broader relevance for other related flaviviruses, such as ZIKV, WNV, and YFV, due to the structural and functional similarities of their capsid proteins [17,21,44,45]. Moreover, DENV C binding to host lipid droplets has been thoroughly studied, resulting in the development of potential peptide-based therapeutic approaches against this and related viruses [21,46,47]. A thorough understanding of DENV C binding to the viral RNA may therefore also enable new therapeutic approaches. With this in mind, it was recently established that key stretches of DENV viral RNA sequence are fundamental for C protein binding [43]. Using such information to design ssDNA analogue proxies allows us to study such interactions while, simultaneously, exploring the possibilities of using those analogues as competitive inhibitors of such interactions in future drug development approaches, against DENV and/or related viruses (including ZIKV, where similar observations were obtained [43]).

## 2. Results

### 2.1 Selection of nucleic acid sequences

Previous experiments [43] mapped the association between the viral genome and DENV C *in vitro*, suggesting that their binding is not uniform. A strong preference was observed for regions involved in long-range intramolecular RNA-RNA interactions. To further characterize the C protein interaction with nucleic acids, viral RNA sequences identified in previous work as crucial for the interaction [43] were selected based on their length, GC content (percentage of nucleotides containing the bases guanine or cytosine relative to the total nucleotide content of a DNA or RNA molecule), predicted secondary structure and location within the RNA genome (Figure S1). Single-stranded DNA sequences, analogous to those RNA sequences, were custom synthesized (Table S1). The rationale for the approach is that they could, on one hand, act as competitive inhibitors of their ssRNA counterparts within the genome and, on the other hand, serve to inform on DENV C binding to nucleic acids.

The binding of ssDNA analogues to DENV C was characterized via different biophysical assays. First, ssDNA sequences with different lengths were analyzed. Three sequences will be shown (C, D and G, with 35, 50 and 75 nucleotides, respectively). To test sequence specificity and the importance of the GC content, scrambled sequences and adenine/thymine-rich (A/T-rich) or cytosine/guanine-rich (G/C-rich) sequences were, respectively, generated and their interaction with DENV C was also analyzed (Table S1).

### 2.2 DENV C-ssDNA interaction experiments

#### 2.2.1 Tryptophan environment modification upon ssDNA interaction

DENV C monomer contains a single tryptophan residue, W69, conserved among *Flavivirus* [15]. It is located at the α4-α4’ interface, on the external side of the α3 core, partially responsible for dimer stabilization [16]. Moreover, it allows the interaction between the α2 and α4 helices [15]. Taking advantage of DENV C intrinsic tryptophan fluorescence, steady-state and time-resolved fluorescence spectroscopies were used to obtain information regarding the tryptophan residue microenvironment, expected to act as a reporter of the protein interaction with the proposed ssDNA ligands. As an example, fluorescence emission spectra of DENV C acquired in the absence and presence of increasing concentrations of three ssDNA sequences with different lengths are shown in Figure S2.

For all sequences, fluorescence intensity (and quantum yield) strongly decreases upon increasing ssDNA concentration (Figure 1A), with a less pronounced reduction when DENV C interacts with the shorter sequence (Figure 1A). This W69 fluorescence quenching, which can go up to approximately 80% in some cases, suggests a significant interaction of DENV C with the studied ssDNA sequences, which might be influenced by their length. Interestingly, the maximum emission wavelength also displays a small blue shift (Table S2) at higher [ssDNA]/[DENV C] ratios. This indicates that W69 undergoes a small transition towards a more hydrophobic microenvironment, possibly due to the interaction with ssDNA.

**Figure 1.**
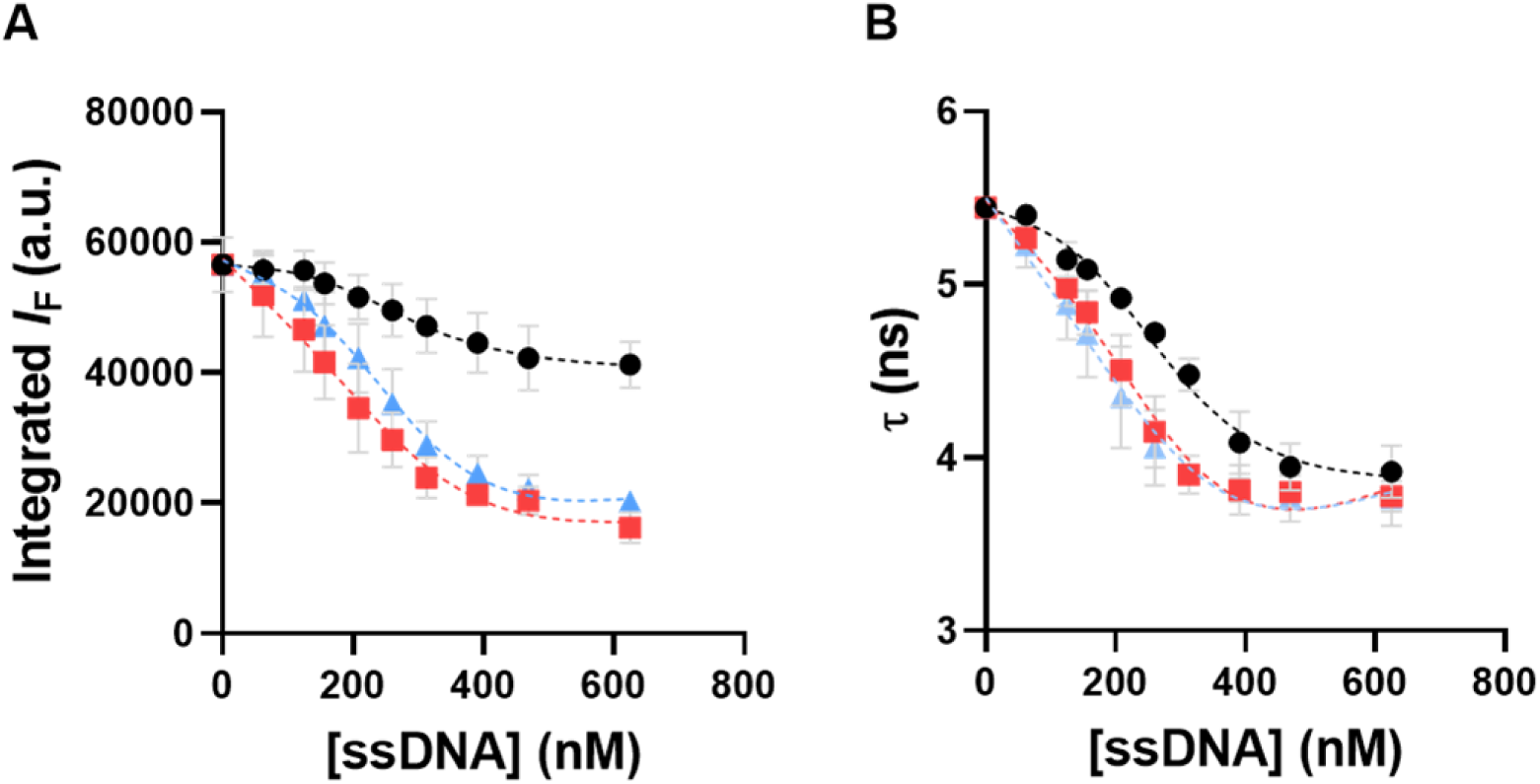
DENV C interaction with ssDNA of different lengths. Integrated fluorescence intensity (**A**) and average fluorescence lifetime (**B**) of DENV C in the presence of increasing ssDNA concentrations. The 35-mer (sequence C), 50-mer (sequence D) and 75-mer (sequence G) sequences are represented by black circles, red squares, and blue triangles, respectively. Initial DENV C concentration was kept at 2 µM for all ssDNA sequences. ssDNA concentration varied from 62.5 nM to 625 nM. Error bars represent standard deviation of at least 2 independent experiments. Dashed lines are guides to the eye.

To further explore this DENV C-ssDNA interaction, time-resolved fluorescence emission measurements were carried out. Fluorescence lifetime values obtained at different ssDNA concentrations are depicted in Figure 1B. As observed in the steady-state measurements, DENV C lifetime strongly decreases with increasing ssDNA concentrations, until it reaches a plateau-like stabilization, suggesting a saturation of the protein with the nucleic acid and, once again, pointing to a significant protein-ssDNA interaction.

To better understand the mechanism behind the observed fluorescence quenching of DENV C by ssDNA, Stern-Volmer plots were built both for steady-state and time-resolved fluorescence data (Figure 2). As a rule, if dynamic quenching occurs, both data sets can be described by linear regressions with positive slope [48]. In this case, the quencher would diffuse to the fluorophore during the lifetime of the excited state, without irreversible changes. If static quenching is the main responsible for the results obtained, the Stern-Volmer plot constructed using time-resolved data will present a slope close to 0, due to the formation of a nonfluorescent ground-state complex between quencher and fluorophore [48]. As depicted in Figure 2, the Stern-Volmer plots either based on fluorescence lifetime (Figure 2B) or steady-state (Figure 2A) data exhibit significant deviations from linearity, indicating deviations from classical dynamic and/or static quenching models. Therefore, conventional analyses are not applicable, and empirical equations are necessary to quantitatively assess and better understand the quenching behavior.

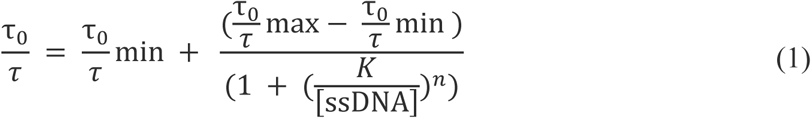

**Figure 2.**
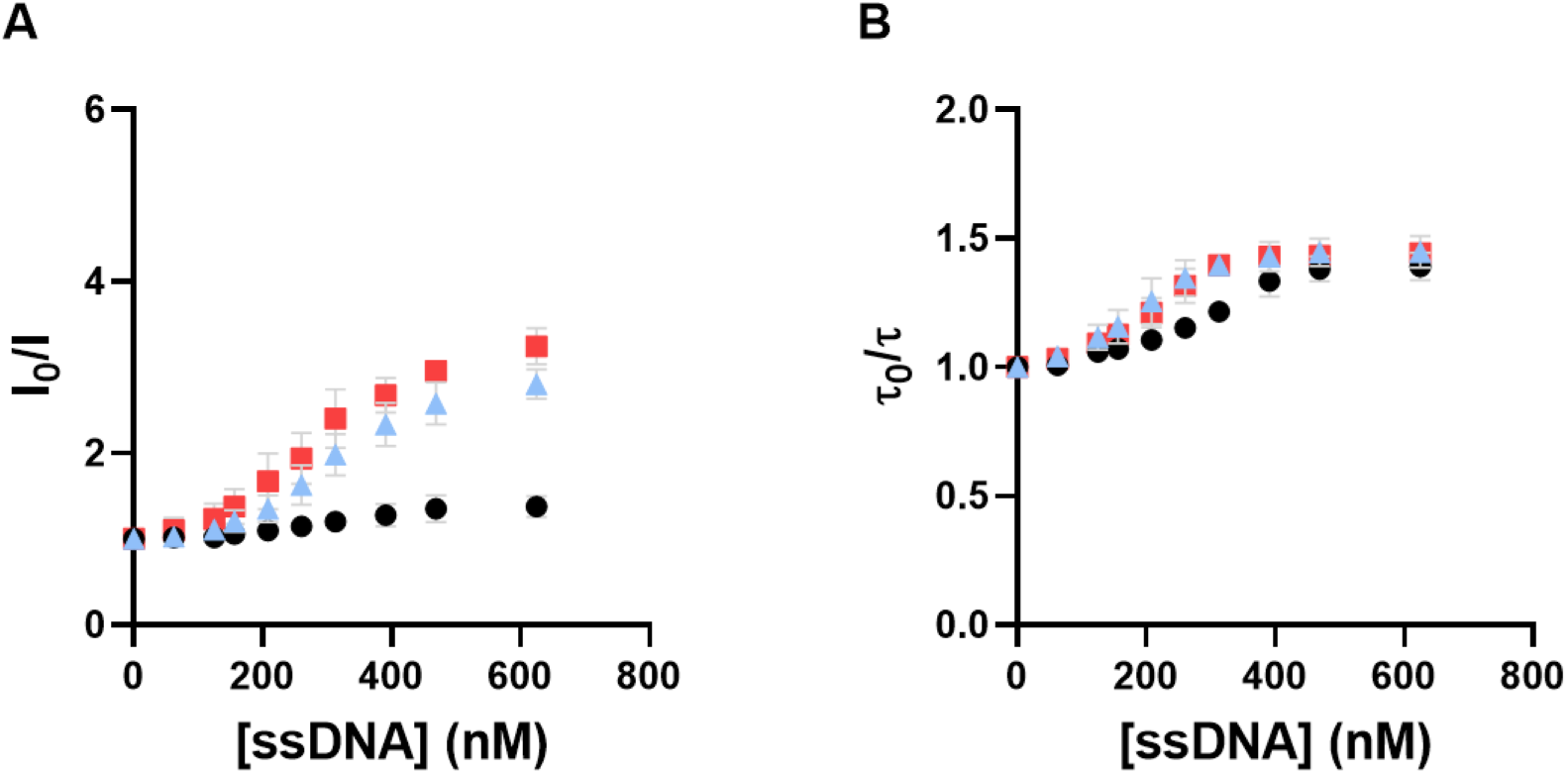
Stern-Volmer plots of DENV C intrinsic tryptophan residue fluorescence quenching by ssDNA sequences with 35 (sequence C, black circles), 50 (sequence D, red squares) and 75 (sequence G, blue triangles) nucleotides. Steady-state (**A**) and time-resolved (**B)** results. Initial DENV C concentration was kept at 2 µM for all ssDNA sequences. ssDNA concentration varied from 62.5 nM to 625 nM. Error bars represent the standard deviation of at least 2 independent experiments. Due to the apparent sigmoidal behavior of steady-state and time-resolved data, a Hill-based equation was fitted to the data [49,50]:

where τ is the average fluorescence lifetime obtained for a given ssDNA concentration, τ_0_ is the average fluorescence lifetime in its absence, 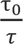 min and 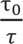 max correspond to the lower and upper limits of 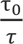, respectively, and *K* is a dissociation constant. The parameter *n* (Hill coefficient) represents the steepness of the curve. The Hill coefficient and *K* values obtained by fitting equation 1 to both steady-state (Figure 3A) and time-resolved data (Figure 3B) are summarized in Table 1.

**Figure 3.**
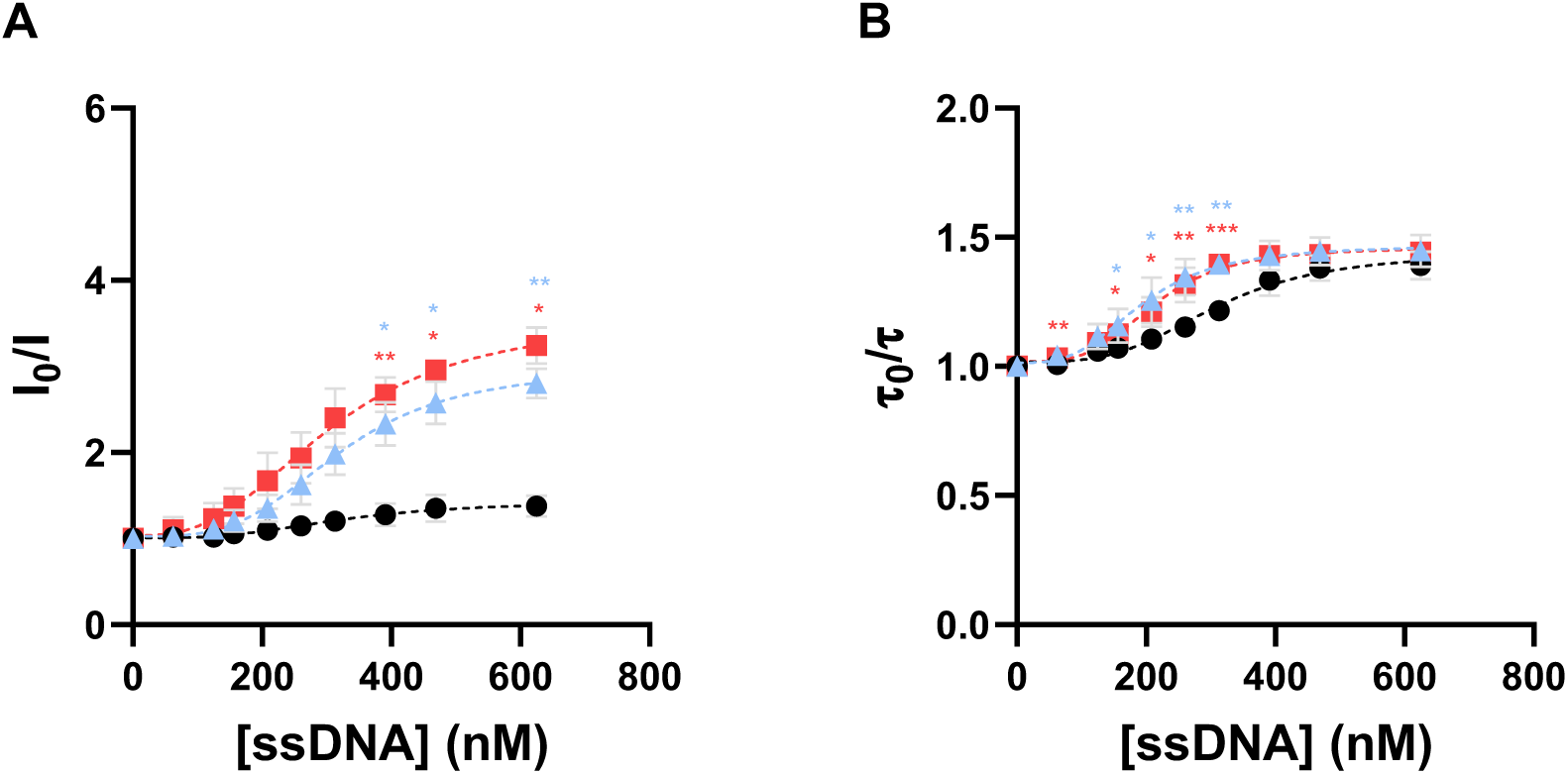
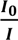 (*A*) *or* 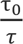 (*B*) variation upon increasing concentrations of ssDNA with 35 (sequence C, black circles), 50 (sequence D, red squares) and 75 (sequence G, blue triangles) nucleotides. Hill equation (1) was fitted to the data (dashed lines). Initial DENV C concentration was kept at 2 µM for all ssDNA sequences. ssDNA concentration varied from 62.5 nM to 625 nM. Error bars represent standard deviation of at least 2 independent experiments. Statistical analysis was performed using the mixed-effects model with Holm-Sidak corrections for multiple comparisons. Asterisk colors correspond to sequence comparisons: red indicates comparisons between sequences C and D, while blue represents comparisons between sequences C and G. *** *p* ≤ 0.001; ** *p* ≤ 0.01; * *p* ≤ 0.05.

**Table 1.**
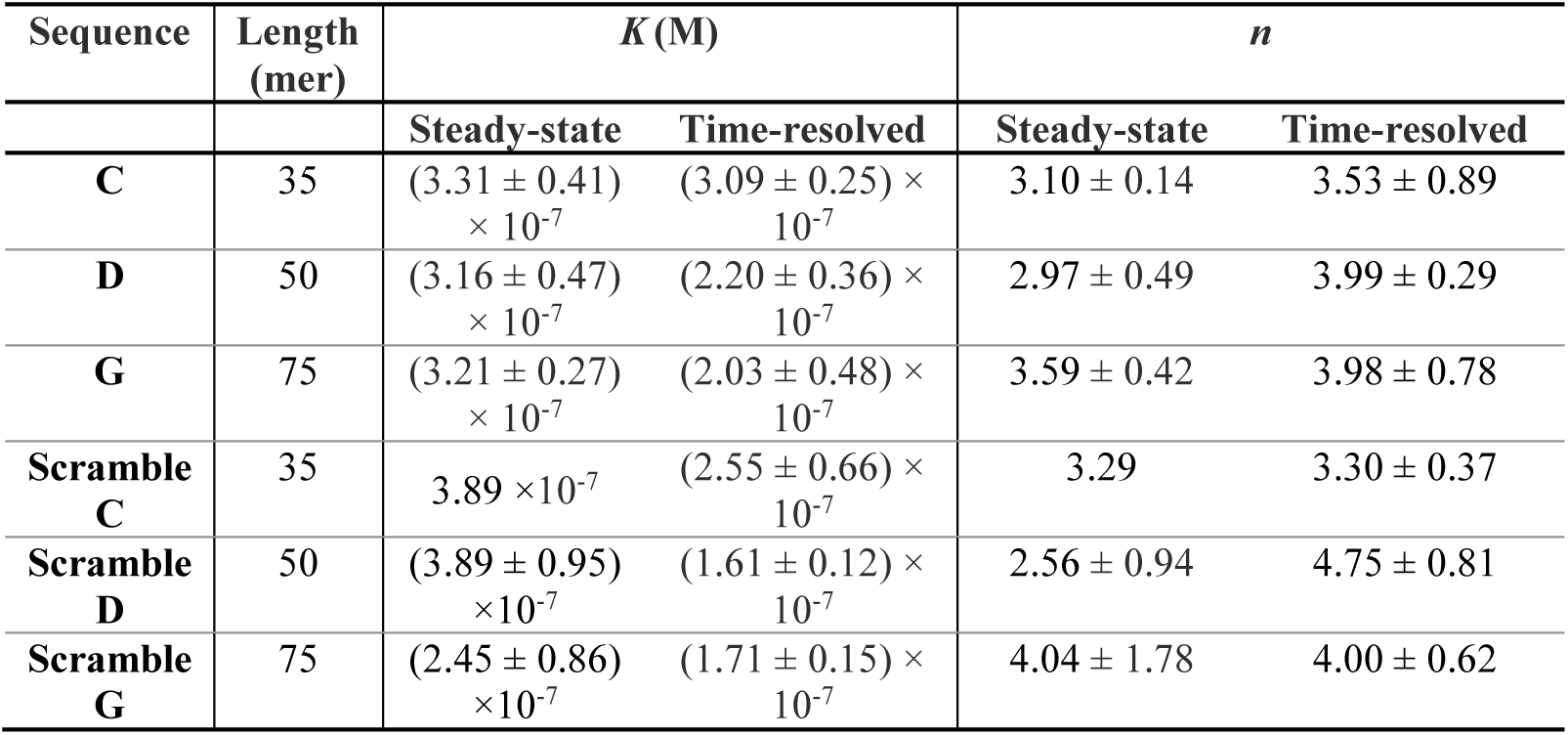
Values of *K* and Hill coefficient (*n*), based on steady-state and time-resolved fluorescence data. Results are expressed as mean *±* standard deviation from at least 2 independent experiments (except for the steady-state data of sequence scramble C).

Both fluorescence intensity (steady-state) and lifetime (time-resolved) variations follow a sigmoidal pattern, indicating that fluorescence changes are likely linked to the cooperative binding of the protein to ssDNA. As shown in Table 1, the Hill coefficient remains consistent across all sequences, suggesting that ssDNA length does not have a major impact in the cooperative nature of the binding. However, despite similar coefficients and lifetime variations (Table 1), the fluorescence quenching efficiency was lower for the shortest ssDNA (Figures 1A and 3A). This suggests that differences in quenching may arise not from variations in binding nature, but rather from variations in fluorophore environment or structural rearrangements within the protein-DNA complex.

#### 2.2.2 Role of sequence specificity in the interaction with DENV C

To test if the nucleotide sequence itself could impact the interaction between DENV C and ssDNA, scrambled sequence DNA molecules based on the previous sequences were randomly generated and analyzed (Figure 4 and Table 1).

**Figure 4.**
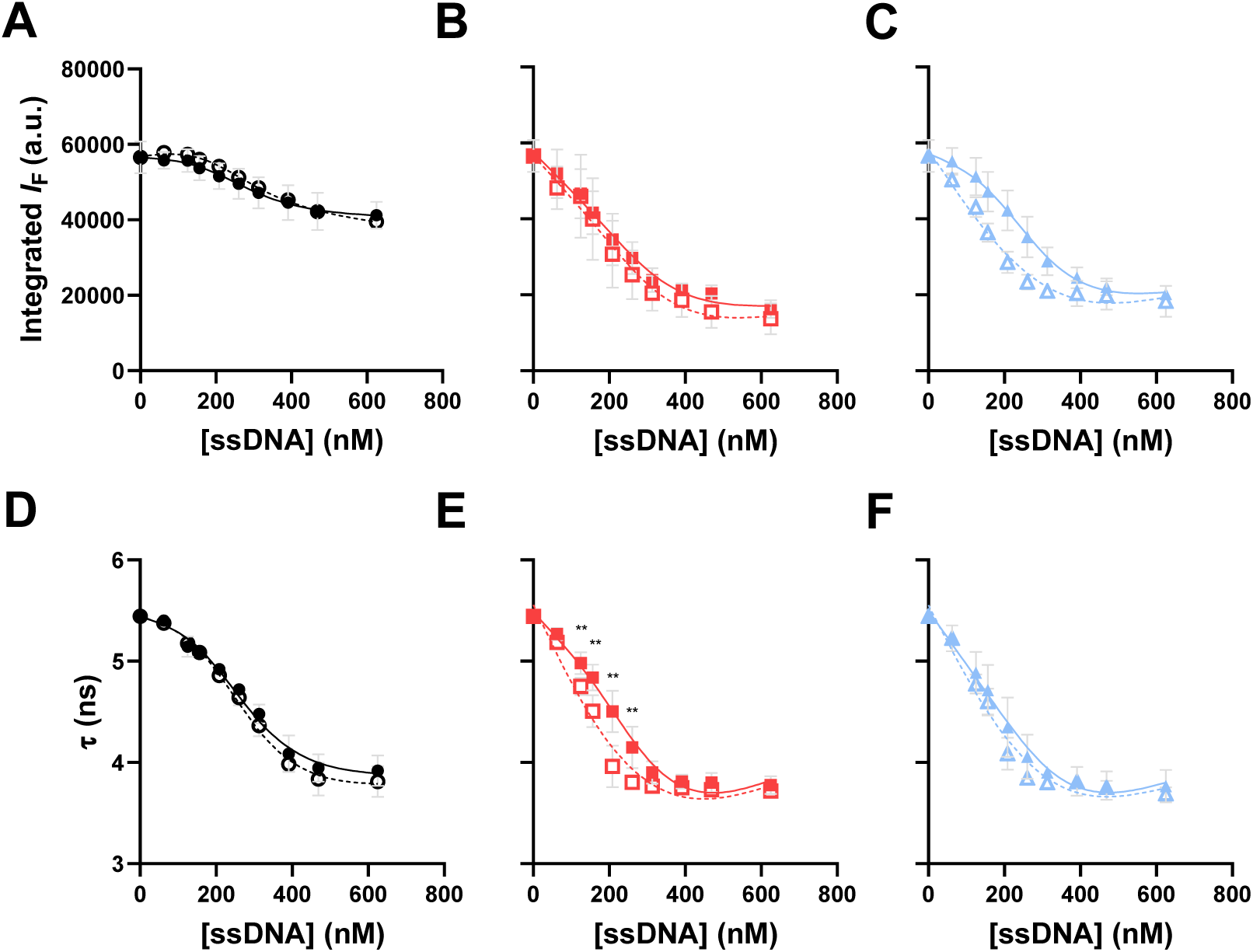
Role of nucleotide sequence in DENV C-ssDNA interaction. Integrated fluorescence intensity (**A-C**) and average fluorescence lifetime (**D-F**) of DENV C in the presence of increasing ssDNA concentrations. DENV C interaction with sequences with 35 (**A, D**), 50 (**B, E**), and 75 nucleotides (**C, F**). Closed symbols represent the viral-based sequences and open circles their respective scrambled sequences. Initial DENV C concentration was kept at 2 µM for all ssDNA sequences. ssDNA concentration varied from 62.5 nM to 625 nM. Error bars represent standard deviation of at least 2 independent experiments. In **A-F**, solid (viral-based sequences) and dashed (scrambles) lines are guides to the eye. Statistical analysis of the data was performed using a multiple comparison t-test with the Holm-Sidak correction. ** *p* ≤ 0.01.

As previously observed, both fluorescence intensity and average lifetime decreased upon increasing ssDNA concentration (Figure 4). Slight changes were observed, but most were not statistically significant. This could indicate that nucleotide sequence may not strongly interfere in the interaction with DENV C. As for the viral-based sequences, Stern-Volmer plots were constructed for both steady-state and time-resolved data (Figure 5). The scrambles exhibited similar sigmoidal behavior, and fitting to the Hill equation yielded parameter values comparable to those obtained for the original counterparts (Table 1).

**Figure 5.**
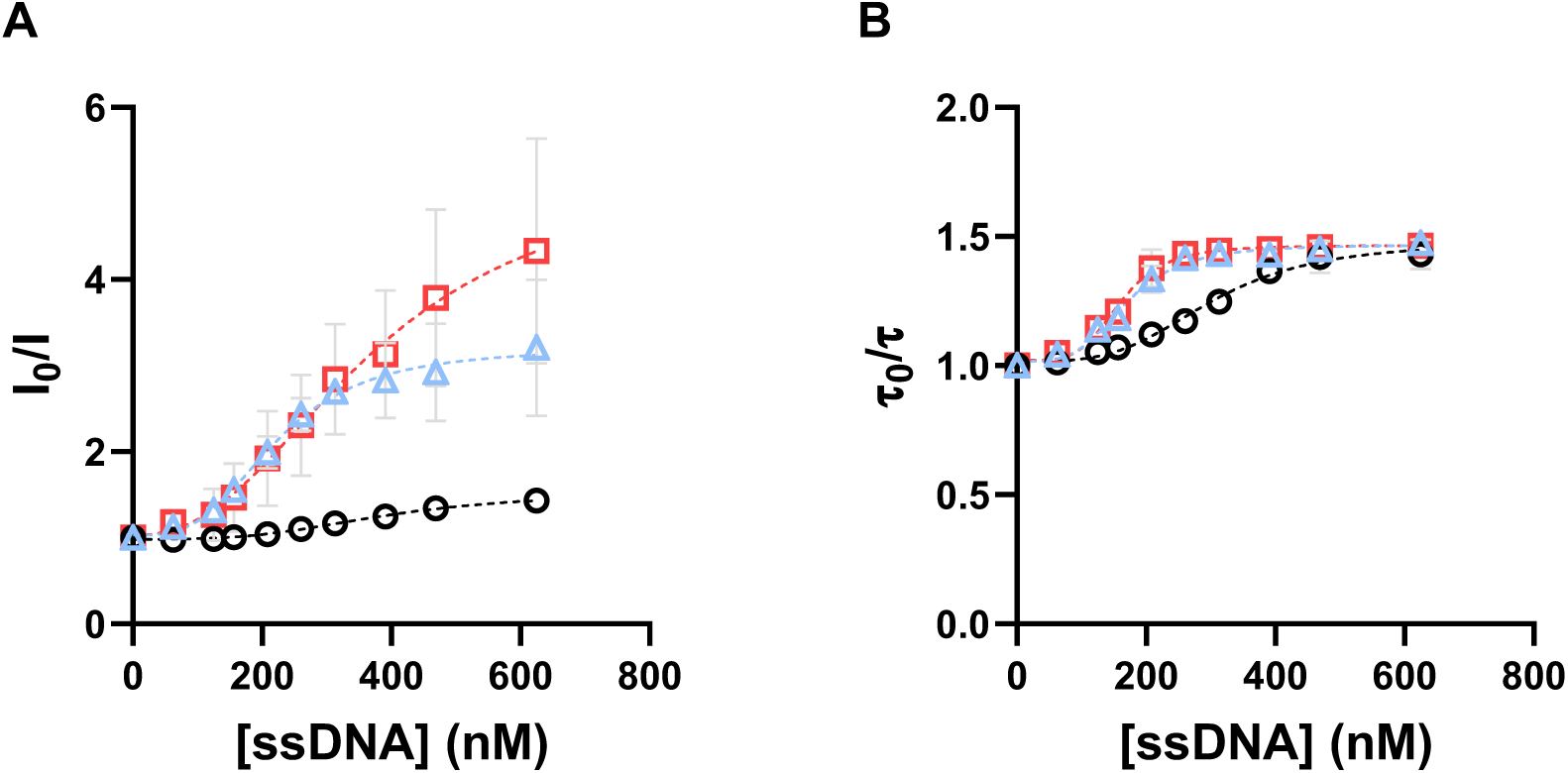
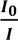 (*A*) *or* 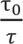(*B*) variation with increasing concentration of ssDNA with 35 (sequence scramble C, open black circles), 50 (sequence scramble D, open red squares) and 75 (sequence scramble G, open blue triangles) nucleotides. Equation 1 was fitted to the data (dashed lines). Initial DENV C concentration was kept at 2 µM for all ssDNA sequences. ssDNA concentration varied from 62.5 nM to 625 nM. Error bars represent the standard deviation of at least 2 measurements.

Overall, fluorescence intensity and lifetime measurements display similar sigmoidal variations for both the original sequences (Figure 3) and their scrambled counterparts (Figure 5), suggesting that sequence scrambling does not significantly impact the overall binding behavior. Furthermore, Hill equation fitting reveals comparable *K* and *n* values across all sequences, indicating that the binding cooperative nature is not strongly dependent on a specific nucleotide sequence. Moreover, differences observed across different lengths also follow the same pattern as those seen for the viral-based sequences. These results suggest that DENV C primarily recognizes general structural features of ssDNA, such as charge distribution, rather than relying on strictly sequence-specific interactions. The similarity in quenching behavior between original and scrambled sequences further supports the idea that the fluorophore environment is affected similarly in both cases, reinforcing the hypothesis that the binding mechanism is not sequence-specific.

#### 2.2.3 Importance of GC content for DENV C-ssDNA interaction

In addition to length and nucleotide sequence, GC content was also analyzed due to its known importance on the physicochemical properties of DNA. The GC content of the original sequences ranged from 43 to 56% (Table S1). To better understand its impact, sequences of 35 or 75 nucleotides with either 0% GC content (A/T-rich) or 100% GC content (G/C-rich) were synthesized and analyzed by time-resolved fluorescence spectroscopy (Figure 6).

**Figure 6.**
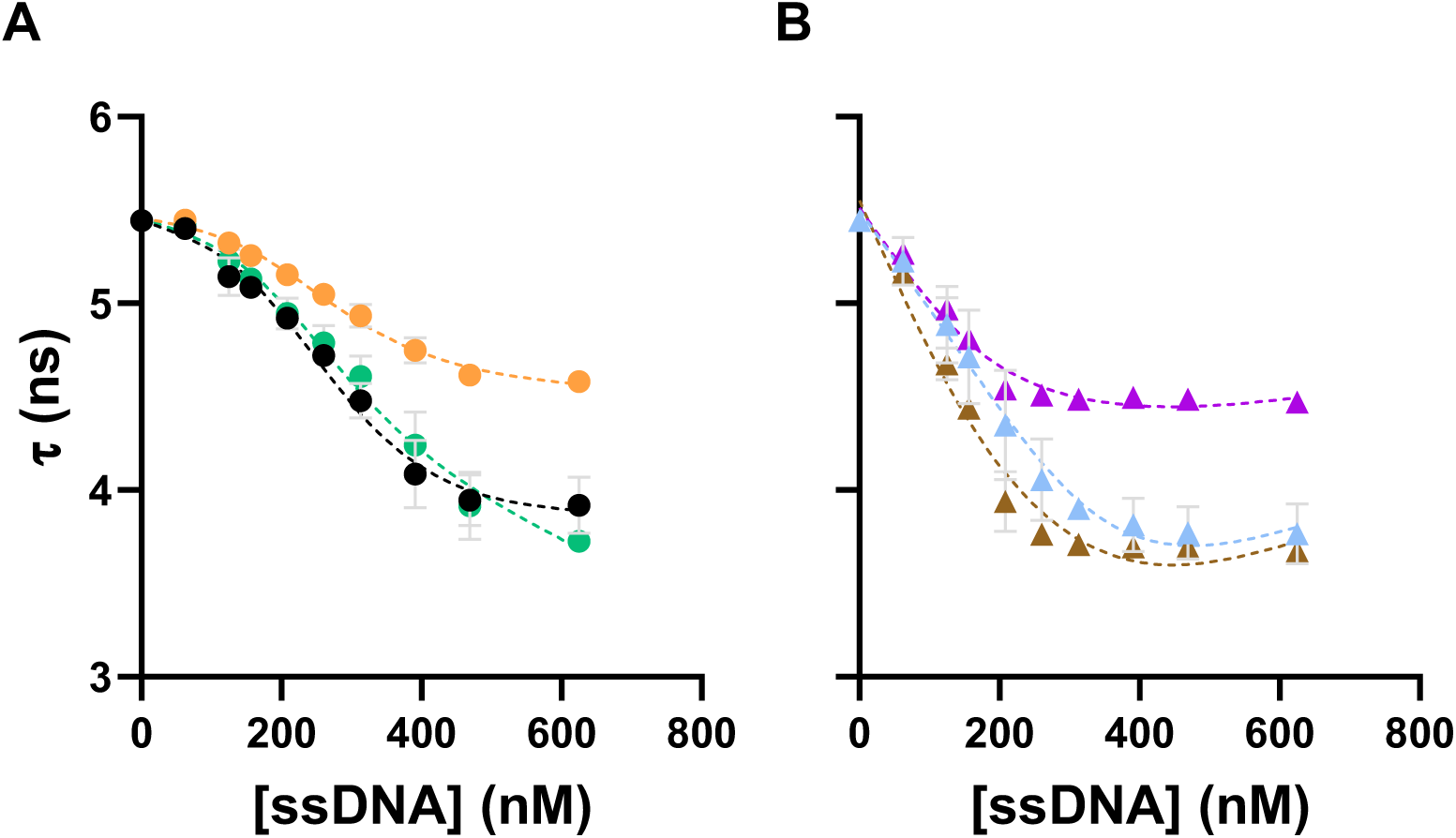
Effect of GC content on DENV C-ssDNA interaction. Average fluorescence lifetime of DENV C in the presence of increasing ssDNA concentrations. DENV C interaction with A/T-rich (orange circles) and G/C-rich (green circles) sequences with 35 nucleotides (**A**), and A/T-rich (purple triangle) and G/C-rich (brown triangles) sequences with 75 nucleotides (**B**) are compared with the sequences with 35 nucleotides (sequence C, black circles) and 75 nucleotides (sequence G, blue triangles), which were included in the respective panel. Initial DENV C concentration was kept at 2 µM for all ssDNA sequences. ssDNA concentration varied from 62.5 nM to 625 nM. Error bars represent standard deviation of at least 2 independent experiments. Lines are guides to the eye.

As previously observed, the average fluorescence lifetime decreased upon increasing ssDNA concentrations (Figure 6). However, a difference is noted between sequences with 100% and 0% GC content. This difference is particularly evident considering the length, where the shorter sequences (Figure 6A) do not even reach a plateau-like stabilization, as observed for the longer sequences (Figure 6B). These findings indicate that GC content may influence the interaction between ssDNA and DENV C, potentially affecting binding affinity or structural properties of the complex. While data for the 35-nucleotide GC-rich sequence may be ambiguous, the overall quenching trend remained consistent with other sequences. These differences may indicate subtle variations in binding behavior or fluorescence environment, deserving further investigation.

#### 2.2.4 Conformational and dynamic insights into DENV C-ssDNA interaction

To further investigate potential structural differences in protein-ssDNA interactions, fluorescence anisotropy and dynamic light scattering (DLS) experiments were performed for some of the ssDNA sequences. These measurements aimed to provide additional insights into binding-induced conformational changes and complex formation. Although anisotropy and DLS data are not available for all sequences, as all gave similar results in the previous experiments, the results obtained here can still help to elucidate the overall structural properties of the protein-DNA complex.

Fluorescence anisotropy measurements (Figure 7) revealed a progressive increase in fluorescence anisotropy upon increasing ssDNA concentration, indicating a reduction in the rotational mobility of the fluorophore.

**Figure 7.**
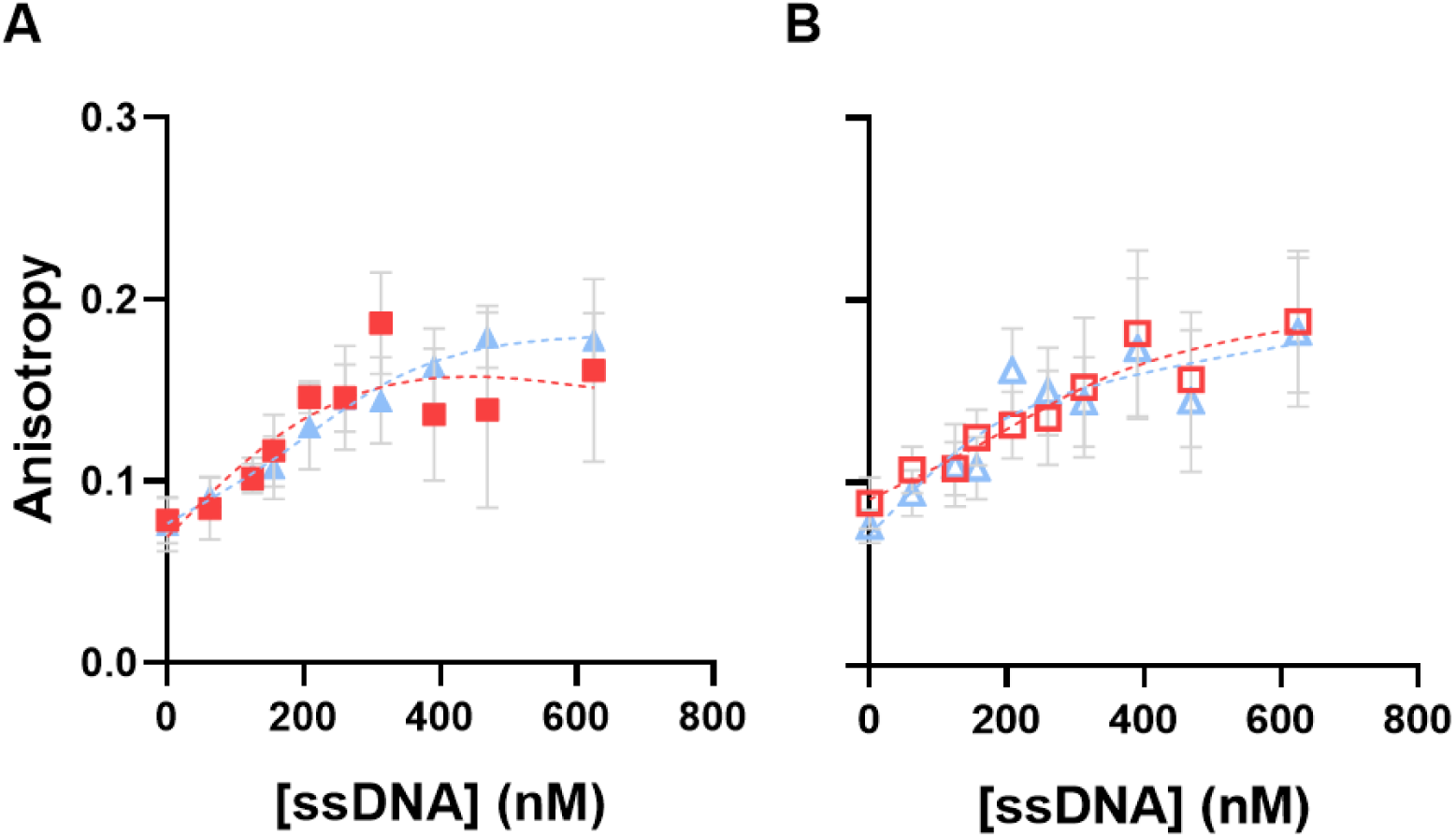
Fluorescence anisotropy variation with increasing concentrations of ssDNA. DENV C interaction with sequences with 50 (sequence D, red squares) and 75 (sequence G, blue triangles) nucleotides (**A**), and their respective scramble counterparts (**B**), with scramble D and scramble G represented by the open red squares and open blue triangles, respectively. Initial DENV C concentration was kept at 2 µM for all ssDNA sequences. ssDNA concentration varied from 62.5 nM to 625 nM. Error bars represent the standard deviation of at least 2 experiments. Dashed lines are guides to the eye.

Furthermore, measurements showed a similar tendency for increasing anisotropy values across all studied sequences (middle and long, original and scrambled), indicating that protein binding leads to a comparable reduction in rotational mobility regardless of ssDNA sequence composition. As previously stated, this suggests that sequence specificity does not significantly alter the overall structural mobility of the protein-ssDNA complex. Overall, this indicates that upon binding, the protein-DNA complex exhibits restricted mobility, likely due to an increase in supramolecular assembly size and/or structural stabilization of the bound state.

While fluorescence anisotropy indicates a reduction in rotational mobility of the complex, it does not provide direct information about changes in the hydrodynamic properties. To address this, DLS measurements were performed to assess the size alterations associated to DENV C-ssDNA binding. Considering that both the 50 and 75 nucleotides viral-based sequences did not show major differences to their counterparts, only one of the original sequences and two scrambled versions were used here. The size distribution profile of DENV C without ligand shows a hydrodynamic diameter of approximately 7 nm (data not shown), as previously observed [32]. The changes in the z-average values with increasing ssDNA concentration (Figure 8) suggest that protein-ssDNA interaction leads to the formation of larger complexes, supporting binding-induced complex formation or structural rearrangements.

**Figure 8.**
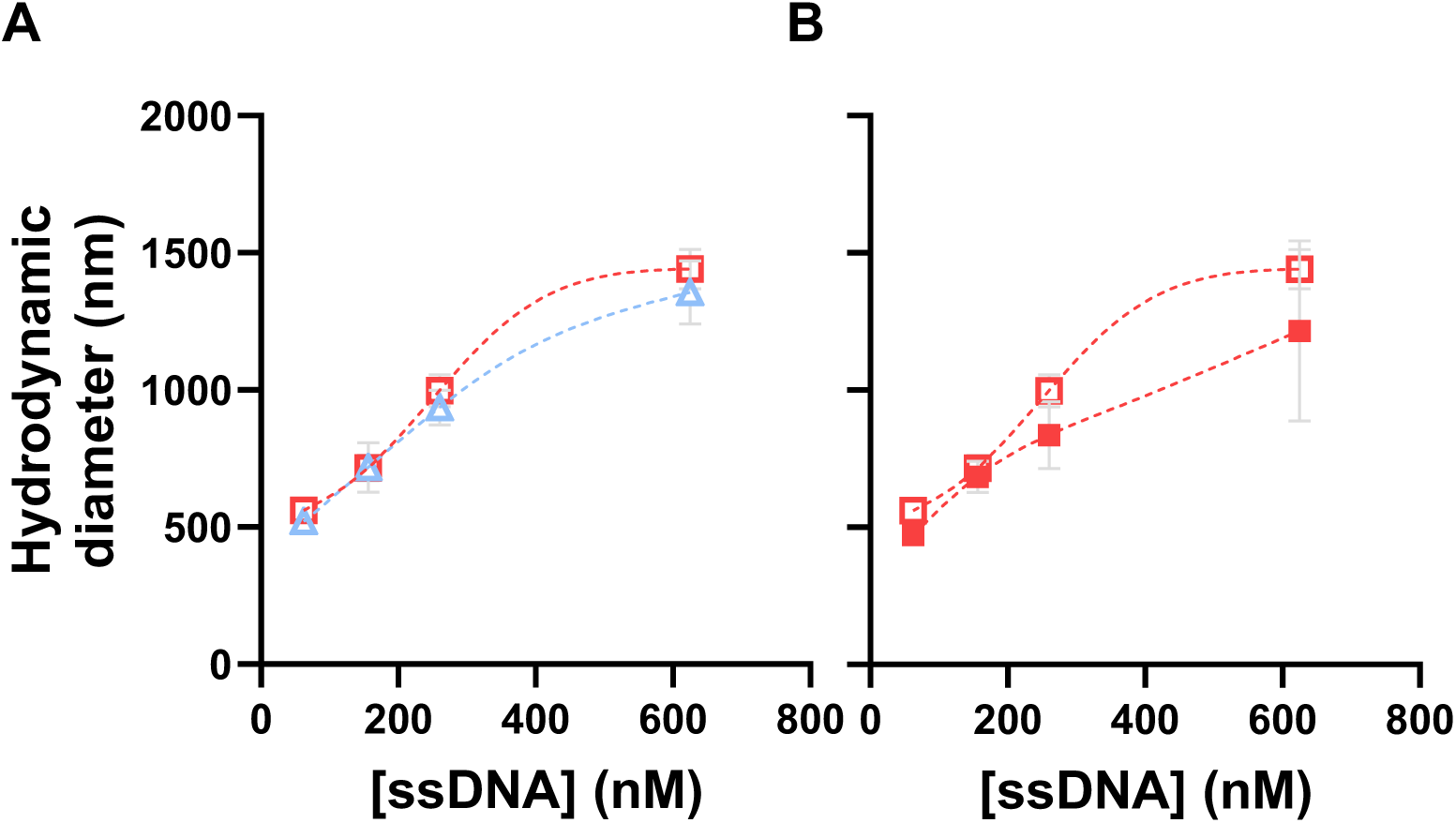
Dynamic light scattering analysis of DENV C-ssDNA complex formation. The z-averaged hydrodynamic diameter is plotted against increasing ssDNA concentration. Comparison of two sequences with different lengths, sequence scramble D, with 50 nucleotides, and sequence scramble G, with 75 nucleotides, represented by the open red squares and blue triangles, respectively **(A)**. Comparison between the original sequence (sequence D, closed red squares) and its scrambled counterpart (sequence scrambled D, red open squares) **(B)**. Initial DENV C concentration was kept at 2 µM for all ssDNA sequences. ssDNA concentration varied from 62.5 nM to 625 nM. Error bars represent the standard deviation of at least two independent experiments. Dashed lines are guides to the eye.

The polydispersity index also increased upon increasing ssDNA concentration, reaching approximately 0.3. This indicates increased sample heterogeneity, likely due to complex formation or oligomerization. The intensity percent distribution (Figure 9) further supports this observation, showing a shift in the main peak towards larger diameters upon increasing ssDNA concentration. The presence of broader peaks at higher ssDNA concentrations indicates the coexistence of multiple species, with broad size distribution, rather than a single homogeneous complex.

**Figure 9.**
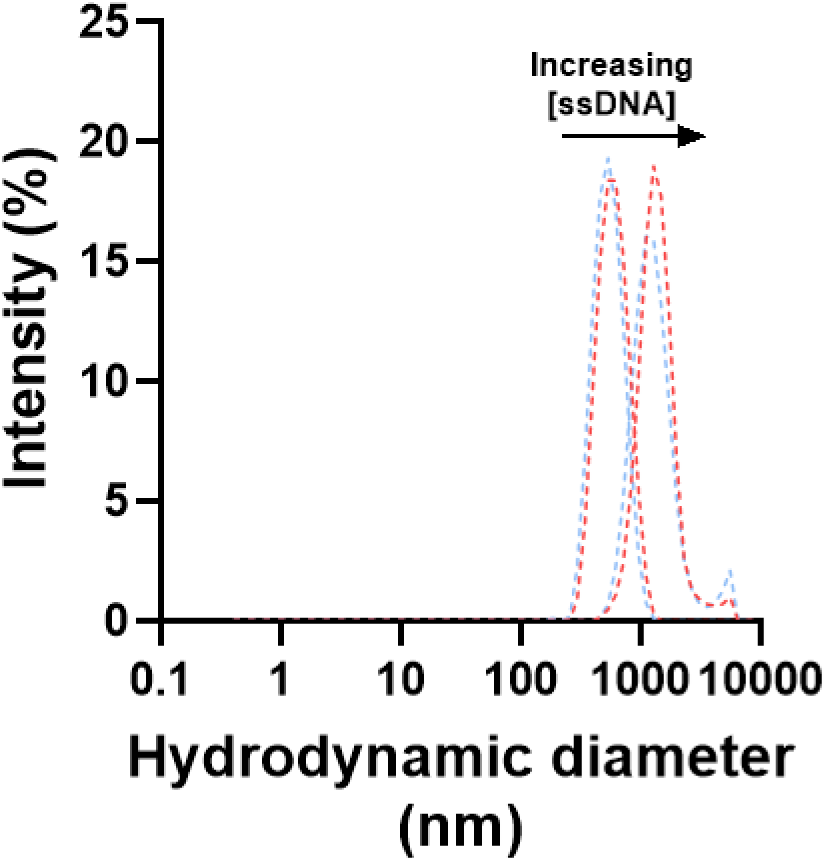
Intensity-weighted size distribution profiles of DENV C-ssDNA complexes in the presence of two ssDNA concentrations. A sequence with 50 nucleotides (sequence scramble D) and 75 nucleotides (sequence scramble G) are represented by the red and blue dashed lines, respectively. Initial DENV C concentration was kept at 2 µM for both ssDNA sequences. Two ssDNA concentrations were tested: 62.5 nM and 625 nM. Peaks at smaller hydrodynamic diameters (left) correspond to the lower ssDNA concentration, while peaks at larger hydrodynamic diameters (right) correspond to the higher ssDNA concentration. The shift toward larger diameters at higher ssDNA concentration demonstrates the formation of higher-order complexes.

The similar tendency between different ssDNA lengths (Figures 8A and 9) and original *vs.* scrambled version (Figure 8B) indicates that all follow a comparable binding pattern, suggesting that, among the analyzed sequences, neither sequence specificity nor length strongly impact complex size, under these conditions. Nevertheless, it cannot be fully excluded a potential length-dependent effect. Furthermore, the possible plateau at higher concentrations suggests that binding saturation may be reached.

### 2.3 DENV C secondary structure changes upon ssDNA interaction

Circular dichroism (CD) spectroscopy was employed to monitor changes occurring on the protein secondary structure upon interacting with ssDNA (Figure 10). Upon titrating ssDNA with DENV C, it is possible to observe that as the [DENV C]/[ssDNA] ratio decreases (*i.e.*, as DENV C concentration is lower), there is a significant modification of the protein secondary structure, with an apparent loss of the α-helical content, an effect clearly seen on the longer sequence (Figure 10C). These changes are quite surprising, since the protein was shown to be stable, even upon increasing the temperature [17].

**Figure 10.**
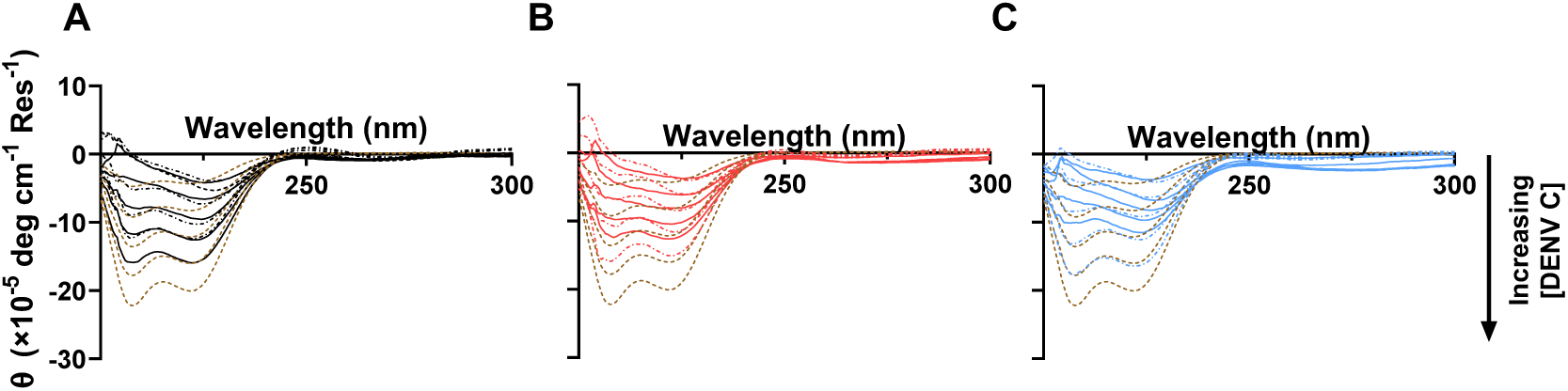
CD spectra of DENV C upon interacting with ssDNA sequences with 35 (sequence C) (A), 50 (sequence D) (B) and 75 nucleotides (sequence G) (C). The original sequence is depicted in solid lines and the respective scramble in dash-dotted lines. Dashed brown spectra represent increasing protein concentration in the absence of ssDNA. In the experiments with ssDNA, its concentration was kept at 125 nM. Protein was sucessively added from 0.4 µM up to 2 µM, corresponding to each spectrum.

The ellipticity at 222 nm was plotted as a function of [DENV C]/[ssDNA] ratio, fitting a linear regression to the data (Figure 11).

**Figure 11.**
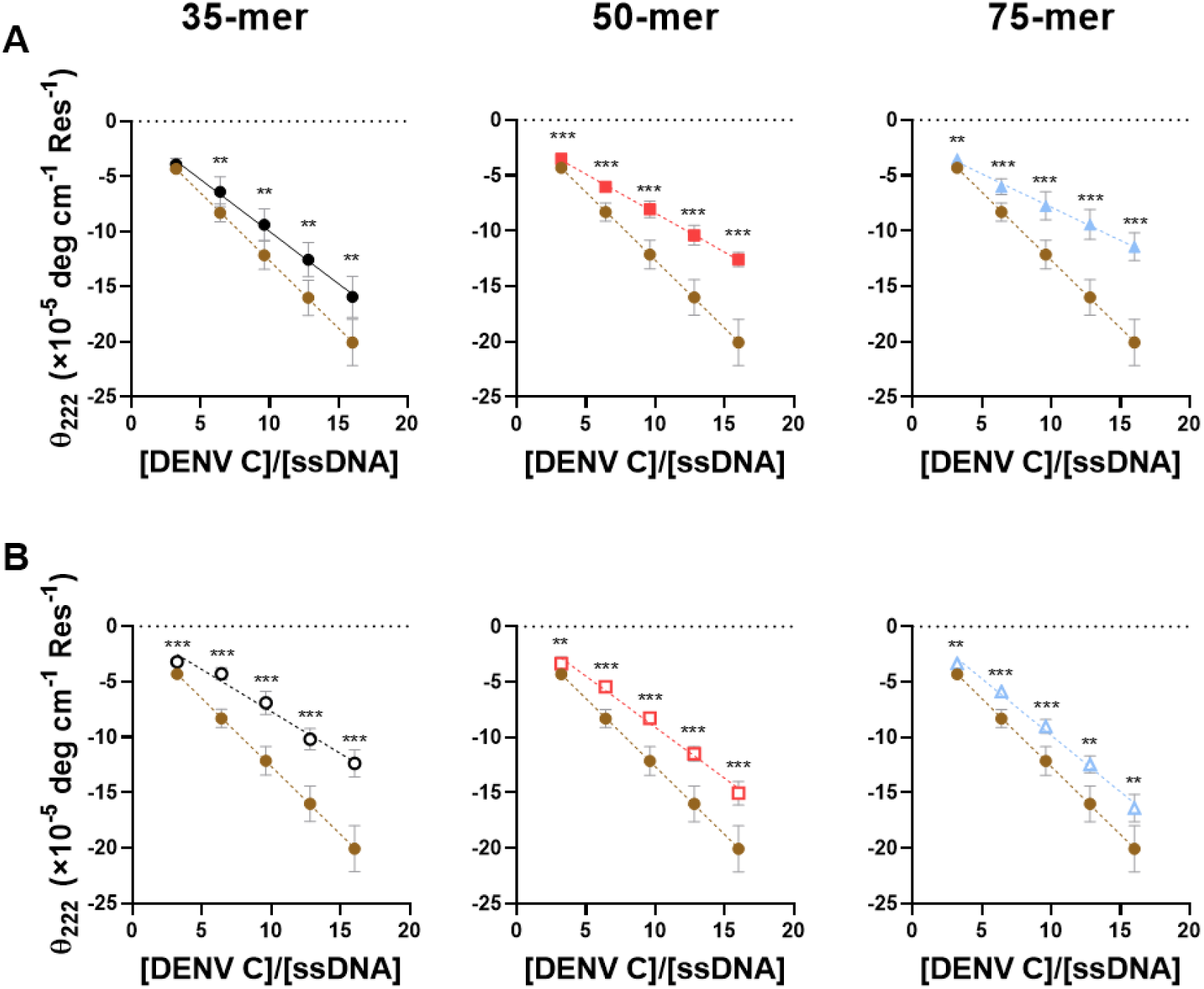
Ellipticity at 222 nm as a function of [DENV C]/[ssDNA] ratio for the original sequences with 35 (sequence C, black circles), 50 (sequence D, red squares) and 75 (sequence G, blue triangles) nucleotides **(A)**, and their respective scrambles with 35 (open black circles), 50 (open red squares) and 75 (open blue triangles) nucleotides **(B)**. Brown solid circles correspond to the control experiment (successive protein additions in the absence of ssDNA). Linear regression fitting was applied to the data (dashed lines). In the experiments with ssDNA, its concentration was kept at 125 nM. Protein was sucessively added from 0.4 µM up to 2 µM. Error bars represent the standard deviation of at least 3 experiments. Statisical significance was assessed by Mann-Whitney U-test. ** *p* ≤ 0.01; *** *p* ≤ 0.001.

The slopes obtained, normalized to control without ssDNA, are presented in Figure 12.

**Figure 12.**
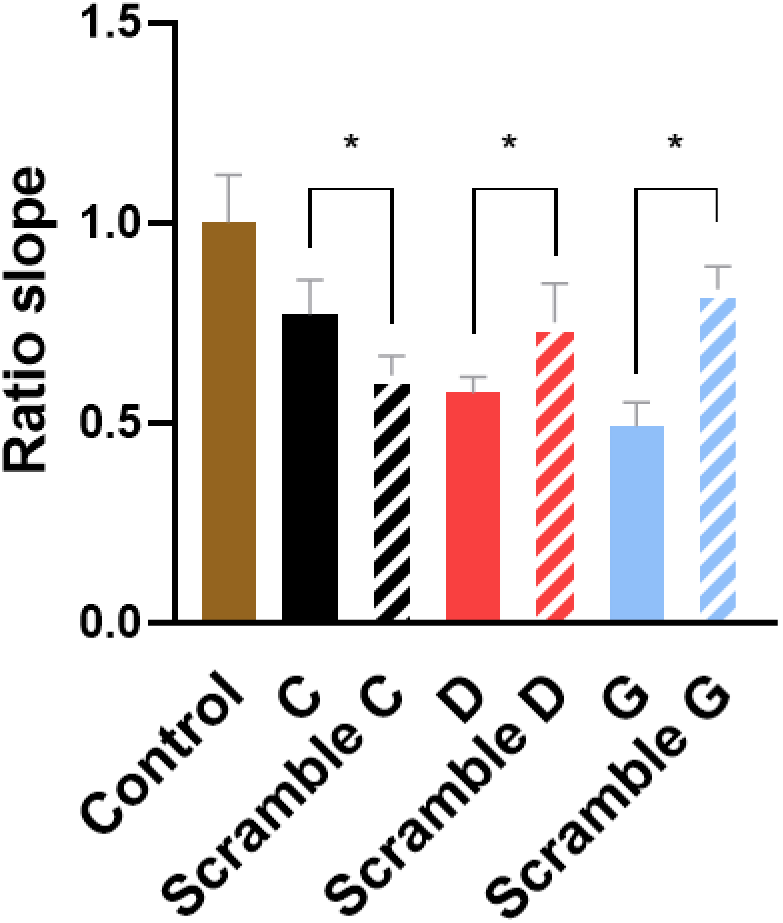
Ratio of slopes of θ_222_ *vs.* [DENV C]/[ssDNA] normalized to the control (protein without ssDNA). Slopes were obtained by fitting a linear regression to the data from ellipticity at 222 nm as a function of [DENV C]/[ssDNA] ratio. Error bars represent the standard deviation of at least 3 experiments. Mann-Whitney U-test was used to assess statistical differences between the sequence and its respective scramble. * *p* = 0.03.

For the original sequences, a relation between the length of the sequence and the steepness of the slope is observed, with longer sequences causing a lower slope (Figure 11A). These results corroborate the hypothesis that possibly there is some length-dependence on the observed changes. Moreover, scrambled sequences with medium and longer lengths seem to cause less extensive conformational changes (Figures 10, 11 and 12), comparing to their original counterparts.

## 3. Discussion

Even though capsid assembly and viral genome packaging are essential steps of the viral life cycle [5,13–15,17,20,21,23,24,46,51], they are still poorly understood. In the case of *Flaviviridae*, the exact number of capsid proteins per viral particle is unknown, making it an exclusion criterion for studies focused on understanding viral packaging strategies based on protein structure and charge [52]. Nevertheless, it is likely that the ratio RNA:C protein is between 1:100 to 1:250, as suggested by recent findings [43]. Supporting this, another study speculated that approximately 180 dimers are required to stabilize the capsid [32].

In simple terms, specific interactions between viral genome and the capsid protein are expected to be a factor promoting RNA packaging selectivity, even though other factors are also involved [53]. While packaging signals within the viral genome have already been identified for some viruses, none have been found for flaviviruses [53]. A proposed mechanism is that selective packaging is mediated by RNA replication-coupled assembly within the assembly sites [53,54]. Another hypothesis suggests that interactions between C protein and viral RNA are unspecific, and electrostatics mediated [54,55].

Recent studies reported that DENV nucleocapsid assembly requires neutralization of the C protein positively charged residues (such as R85 and K86 in the α4-α4’ region) through interactions with nucleic acids or negatively charged surfaces. This process must be tightly coordinated to prevent unspecific aggregation [26]. Nonetheless, C protein-viral genome interactions occur during viral packaging and assembly processes. These interactions promote structural changes in viral genome, as well as in the protein, in order to form a stable complex. It is known that single-stranded DNA or RNA lack a defined structure, enabling electrostatic interactions between the phosphate group and positively charged residues, but also π-π stacking interactions between the free bases and the aromatic residues chains, being the strength of the latter likely responsible for sequence specificity [56]. This may explain why changes in the fluorescence parameters were observed upon interaction with ssDNA. In fact, other studies assessing the interaction of polyelectrolytes with DNA demonstrated that ssDNA is a good quencher, even better than double-stranded DNA [57]. In terms of fluorescence intensity, the signal decreases upon increasing ssDNA concentrations (Figures S2 and 1A), which may be justified by quenching of W69 either by the ssDNA itself and/or, indirectly, by DENV C amino acid residues affected by the interaction. Lysine, glutamine, asparagine, glutamic and aspartic acid may act as tryptophan fluorescence quenchers [58], being all these amino acid residues present in DENV C primary structure. More specifically, lysine residues are the most abundant in DENV C and are in the proximity of W69 (Figure S3), supporting that hypothesis.

Interestingly, the classical Stern-Volmer analysis did not adequately describe either the steady-state or the time-resolved data, suggesting a more complex mechanism underlying DENV C-ssDNA interaction. Therefore, an empirical model was applied in order to better describe the mechanism. Given the sigmoidal shape of the plots, a cooperative model was used [49,50]. The *K* values in the nanomolar range suggest a moderate affinity of ssDNA for DENV C, while the positive Hill coefficient indicates positive cooperativity. A similar behavior was already observed for the interaction of polyelectrolytes and DNA, and the quenching mechanism was well modeled in terms of a complexation equilibrium in which a PFP-NR3/DNA aggregate complex is formed, bringing polymer chains into close enough proximity to allow interchain excitation energy migration and quenching within the aggregate [57].

Aside from the clearly lower quenching efficiency observed for the shortest sequence (Figure 1A), only slight differences were found between sequences of different lengths, as well as between viral-based sequences and their scrambled counterparts. Given the crucial role of GC content in determining the physicochemical properties of DNA, we further analyzed its importance in the context of DENV C-ssDNA interaction (Figure 6). W69 fluorescence lifetime variation was significantly affected by sequences rich in guanines and cytosines. This might be explained by at least two different hypothesis: *i*) a higher affinity of DENV C for GC-rich sequences; or, *ii*) a preference for certain structures that are prone to occur in GC-rich sequences. Guanines were already demonstrated to have a higher quencher efficiency compared to the other bases [57].

Overall, these results suggest that cooperative binding occurs similarly, regardless of sequence length or specificity of the sequences and, that DENV C may primarily recognize general features of ssDNA rather than specific sequences. GC content may be an important factor in DENV C-ssDNA interactions, potentially influencing the affinity. However, further studies are needed to confirm this hypothesis.

Changes in both protein and DNA secondary structures are known to occur upon their interaction [59]. Here, CD analysis revealed a large and unexpected change in DENV C secondary structure, characterized by an apparent loss of α-helical content, as well as a decreased signal (Figure 10). Structural changes have been reported for other viral proteins, such as the envelope glycoprotein complex of human immunodeficiency virus type 1 (HIV-1) [60]. These results suggest that both sequence length and specificity may play a role in the observed structural changes, as DENV C secondary structure was not equally affected by all sequences. For instance, the longest scrambled sequence did not induce an effect on DENV C secondary structure as pronounced as its original counterpart (Figures 10, 11 and 12). Taken together, a picture starts to emerge in which pure electrostatic-driven events should not be held solely accountable for the level of observed changes.

To further investigate binding-induced changes and complex formation, fluorescence anisotropy and DLS experiments were performed. The observed anisotropy increase (Figure 7) is consistent with a model in which ssDNA binding induces conformational changes in the protein, leading to a more rigid and structured complex. Notably, the magnitude of anisotropy change did not vary between different ssDNA sequences, suggesting that it is likely that neither length nor sequence composition significantly influences the degree of conformational restriction. This finding aligns with the steady-state and time-resolved fluorescence results. A similar increase in anisotropy values has been previously reported using fluorescein-labelled DENV C and oligonucleotides smaller than those used in this study [26]. Interestingly, in that case an oligonucleotide-length dependence was observed, although under different experimental conditions [26].

DLS measurements further revealed a significant increase in hydrodynamic size upon ssDNA binding, indicative of larger complex formation (Figures 8 and 9). This observation raises the possibility that these interactions could lead to a phase-separated state, as observed in other nucleic acid-binding proteins [61]. Additionally, it suggests that DENV C-ssDNA interactions may promote an initial nucleation event, which could serve as a precursor for larger complex formation. In biomolecular condensation, nucleation often guides the transition from soluble to more stable complexes [62]. Since protein-RNA aggregation is tightly regulated to prevent aberrant clustering [63], it is possible that DENV C undergoes a controlled nucleation process. This process may enable a phase transition, while preventing aberrant aggregation, potentially resulting in the formation of biomolecular condensates (BMC) instead of rigid aggregates. Unlike irreversible protein aggregation, liquid-liquid phase separation (LLPS) is a mechanism that can drive the condensation of proteins and nucleic acids into BMC (or membraneless organelles), which display liquid-like properties that enable diffusion within the condensates [64]. This process is known to be facilitated by intrinsically disordered regions in proteins, which are recognized as drivers of LLPS [61,64]. Viruses are known to exploit BMC to facilitate viral replication and genome assembly [64,65]. For instance, in HIV-1, BMC seem to play roles in both viral entry and viral assembly, with phase separation primarily driven by the nucleocapsid domain of the Gag structural protein [64,65]. Similarly, the nucleocapsid protein of SARS-CoV-2 is able to phase separate in response to pH, salt concentration and RNA levels, where low RNA amounts promote phase separation and higher concentrations suppress it [66].

The relevance of LLPS contribution to the observed interaction between DENV C and ssDNA still needs to be explored. However, such phase separation has already been observed under different experimental conditions [30]. In fact, flavivirus capsid proteins are known to localize within BMC, such as nucleoli [67], and the N-terminal region is intrinsically disordered [15,21] (a feature commonly associated with LLPS [67]). Additionally, electrostatic and π-π interactions, which are likely involved in DENV C-ssDNA binding, are known to facilitate phase separation [65]. Such a mechanism could also help to explain the dynamic incorporation of DENV C into the virion, potentially contributing to structural variability. The structural changes observed in DENV C may be relevant in the context of phase separation, potentially contributing to its role in nucleocapsid assembly.

Given the stability of DENV C and the fact that W69 is in a highly stable region and responsible for dimer stabilization [16], CD results were somehow unexpected. Nevertheless, it is possible to hypothesize that such changes are needed to initiate the viral assembly process, with the packaging of the viral genome and its encapsidation proceeding afterwards. Moreover, the fact that the C protein undergoes structural changes might justify why there is no clear information regarding the number of copies of this protein or the structure within the virion. In fact, it is not unlikely that a variable number of DENV C dimers exist within the virion (with some virions with more DENV C molecules than others), as it might adjust itself to better fit and minimize the energy required for stabilization, depending on where and how the mature nucleic acid molecule binds within the virion. Notably, several binding spots have been identified within the viral RNA [43], which may function as alternatives one to another. This could also explain the lack of an ordered and easy to picture virion internal structure [5,8,10–12,40].

Given the observed interactions between DENV C and ssDNA, it is intriguing to consider whether these findings could be leveraged for therapeutic purposes. One can hypothesize that ssDNA sequences may be employed to impair DENV C crucial interaction with the viral genome. Oligonucleotides-based therapeutics is a thriving field, with fomivirsen (Ionis Pharmaceuticals, Carlsbad, CA, USA) being the first Food and Drug Administration-approved antisense oligonucleotide drug [68]. Fomivirsen is used to treat viral retinitis by inhibiting human cytomegalovirus replication [68]. More recently, a combination therapy of a recombinant protein (based on the envelope and capsid proteins of ZIKV) with an oligodeoxynucleotide has been shown to enhance cellular immune response [69]. This underscores the potential of oligonucleotides, which are being actively explored. Further tests would be needed to fully elucidate the mechanism underlying DENV C-ssDNA interactions and their therapeutic potential. Altogether, the present work sheds light on the structural changes that DENV C undergoes upon interaction with nucleic acids, with potential implications for both viral assembly and antiviral strategies.

## 4. Materials and Methods

### 4.1 DENV C protein production and purification

Recombinant DENV C protein expression and purification were based on previous approaches [17,20]. DENV-2 C protein, strain New Guinea (GenBank database: AAC59275.1) [70], was expressed in *Escherichia coli* (*E. coli*) C41, transformed with DENV-2 C protein gene, cloned into a pET21a plasmid. A single *E. coli* colony was inoculated into 6 mL of lysogeny broth (LB) medium with 100 μg/mL ampicillin, and incubated overnight at 220 rpm, 37 °C. The overnight culture was diluted 1:50 in LB, and incubated at 37 °C and 220 rpm, until the optical density at 600 nm reached 0.8-1.0. Protein expression was induced with 0.5 mM isopropyl β-D-1-thiogalactopyranoside, at 20 °C, 200 rpm, up to 16 h. The culture was pelleted by centrifuging at 7000 × *g* for 1 h, at 4 °C. Supernatants were discarded and the cell pellet was resuspended in 20 mL of HEPES buffer 25 mM pH 7.4 with 0.2 M NaCl, 1 mM EDTA, 5% (v/v) glycerol and 10 μM protease inhibitor cocktail (Sigma). Cells were lysed by sonication, keeping the suspension on ice, with 10 pulses of 30 s at maximum intensity and pauses of at least 1 min. Salt concentration was adjusted by adding NaCl to the cell suspension to reach a final concentration of 2 M, and incubated at least 1 h, with agitation, at 4 °C. The suspension was centrifuged twice at 16 100 × *g* for 30 min at 4 °C. Supernatants were collected and kept at -20 °C until purification. Before the first chromatography step, the protein extract was diluted 4-fold. Then, it was injected into a 5 mL HiTrap heparin column (HiTrap Heparin HP, Cytiva, Marlborough, MA, USA) coupled to an ÄKTA purifier system (ÄKTA explorer, GE Healthcare, Little Chalfont, UK). DENV C was then eluted by performing a NaCl gradient from 0.2 M to 2 M NaCl. The fractions corresponding to DENV C were collected and injected into a size exclusion column (Superdex 75 10/300, Cytiva). DENV C was eluted in a pH 6.0 buffer with 55 mM KH_2_PO_4_ and 550 mM KCl, and stored at -80 °C.

### 4.2 Protein quality control

The quality of the samples was assessed by several techniques (Figure S4). Protein purity was assessed by a 15% sodium dodecyl sulfatepolyacrylamide gel electrophoresis (SDS-PAGE) and UV-Vis spectrum. CD was used to confirm the secondary structure, as described below. DLS measurements, described ahead, were also conducted to confirm the absence of large protein aggregates. The exact molecular weight was validated via mass spectrometry analysis. Protein concentration was determined from the sample absorbance at 280 nm, using DENV C protein extinction coefficient, 5500 M^-1^cm^-1^, calculated from its primary structure.

### 4.3 Nucleic acids

Regions of the viral genome were previously identified as potential DENV C binding sites [43]. ssDNA sequences analogous to those regions were commercially obtained (Synbio Technologies, Monmouth Junction, NJ, USA). Scramble sequences were generated using the Sequence Scramble online tool (GenScript, Piscataway, NJ, USA) and produced by Synbio Technologies. Lyophilized ssDNA was resuspended in Milli-Q water to a stock concentration of 100 µM.

### 4.4 DENV C-nucleic acids interaction

DENV C stock solutions (in 55 mM KH_2_PO_4_, 550 mM KCl, pH 6.0) were diluted to a final monomer concentration of 2 μM in 50 mM KH_2_PO_4_, 200 mM KCl, pH 6.0. Each ssDNA to be analyzed was successively added from 62.5 nM up to 625 nM, to a solution containing 2 μM of DENV C. Before each measurement, the sample was incubated for 15 min, at room temperature, to ensure that equilibrium was reached. Experiments were performed in 50 mM KH_2_PO_4_, 200 mM KCl, pH 6.0. All conditions were measured independently, with different protein batches. For CD measurements, ssDNA concentration was kept constant (125 nM), while protein concentration varied from 0.4 to 2 μM. Suitable controls (either without ssDNA or protein) were carried out.

### 4.5 Steady-state fluorescence spectroscopy

Steady-state fluorescence intensity measurements were carried out on a Varian Cary Eclipse fluorescence spectrophotometer (Mulgrave, Australia), equipped with a xenon flash lamp, using 0.5 cm × 0.5 cm quartz cuvettes (Hellma Analytics, Müllheim, Germany). Emission spectra were acquired between 300 and 450 nm, with excitation at 270 nm. Excitation and emission bandwidths were set to 5 nm and 20 nm, respectively. All fluorescence emission data were corrected for background contribution, dilution upon ssDNA addition and inner filter effects, as described elsewhere [71].

### 4.6 Time-resolved fluorescence spectroscopy

Fluorescence decays were obtained through the time-correlated single-photon timing technique [17,72]. Measurements were carried out in a Life Spec II equipment with an EPLED-280 pulsed excitation light-emitting diode (Edinburgh Instruments, Livingston, UK). Tryptophan was excited at 275 nm, while emission was acquired at 350 nm. The instrument response function, IRF(t), was obtained using a suspension of nanoscale spherical particles of amorphous silica dispersed in water (Merck Life Sciences, Darmstadt, Germany) and matching excitation and emission at 275 nm. Measurements were performed in 0.5 cm × 0.5 cm quartz cuvettes, at room temperature. Blank samples were also analyzed, and photon counts were negligible. Decays were analyzed using FAST software (Edinburgh Instruments, Livingston, United Kingdom), considering a multi-exponential model:

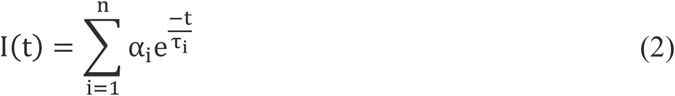

where α_i_ are the pre-exponential factors and τ_i_ the *i*^th^ component of the fluorescence lifetime. The goodness of the fit was evaluated from the experimental χ^2^ value (considered acceptable below 1.4) and global residuals distribution. The intensity average lifetime was described by:

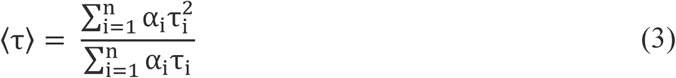

### 4.7 Fluorescence anisotropy

Steady-state fluorescence anisotropy measurements were performed in a Varian Cary Eclipse fluorescence spectrophotometer (Mulgrave, Australia). Fluorescence anisotropy (r) was calculated as [73]:

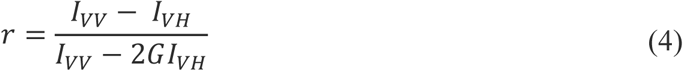

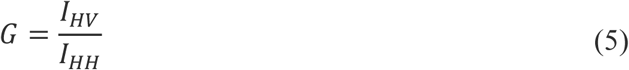

where *Ivv* and *Ivh* are, respectively, the parallel and perpendicular polarized fluorescence intensities measured with vertically polarized excitation light, while *Ihh* and *Ihv* are the parallel and perpendicular polarized fluorescence intensities measured with horizontally polarized excitation light, respectively. The *G* factor is an instrumental correction parameter accounting for the polarization bias in the system. Excitation and emission wavelengths were, respectively, 275 nm and 350 nm. Each anisotropy measure was an average of 6 accumulations. All data were corrected for dilution.

### 4.8 Dynamic light scattering

DLS experiments were carried out at 25 °C on a Malvern Zetasizer Nano ZS (Malvern, UK) with a backscattering detection at 173°, equipped with a He-Ne laser (λ= 632.8 nm), using a quartz cuvette. Samples were left equilibrating for 15 min at 25 °C before each measurement set (10 measurements, each one being the average of 10 runs, with 10 s per run). The scattering intensity weighted data were analyzed using the instrument’s software to determine each sample’s hydrodynamic diameter, size distribution and polydispersity index. The instrument detects time-dependent fluctuations in the intensity of scattered light from particles in suspension. Malvern’s DTS software processes the acquired correlogram (a plot of the correlation function over time) to calculate hydrodynamic diameter [74]. Values were derived from the autocorrelation function using the Cumulants method, which applies a single exponential fit to determine the mean particle size (z-average diameter) and the distribution width (polydispersity index).

### 4.9 Circular dichroism spectroscopy

CD measurements were carried out in a JASCO spectropolarimeter J-815 (Tokyo, Japan), using 1.0 cm or 0.1 cm path length quartz cuvettes, according to the experiment. Spectra were acquired between 200 and 300 nm, with data pitch of 0.5 nm, wavelength sampling velocity of 200 nm/min, data integration time of 1 s and performing 3 accumulations. Experiments were conducted at 25 °C, with temperature controlled by a JASCO PTC-423S/15 Peltier equipment. Spectra were corrected for buffer and ssDNA contribution, and dilution.

### 4.10 Statistical analysis

Results were expressed as mean ± standard deviation of at least two independent experiments. Statistical analyses were performed using different tests, which are specified throughout the results section. *p* ≤ 0.05 was considered as statistically significant. All tests were performed using GraphPad Prism v. 8.0.2 (GraphPad software, Boston, MA, USA).

## Author contribution

**NMS** contributed to conceptualization, methodology, investigation, formal analysis and writing – original draft; **ASM** contributed to conceptualization, methodology, investigation and writing – review & editing; **NEK** contributed to conceptualization, methodology, investigation, formal analysis and writing – review & editing; **MJS** contributed to conceptualization, methodology, investigation, formal analysis and writing – original draft; **RGH** contributed to conceptualization, methodology, investigation and writing – review & editing; **FJE** contributed to methodology and writing – review & editing; **ICM** contributed to conceptualization, methodology, investigation, formal analysis, writing – original draft, funding acquisition, supervision, and project administration; **NCS** contributed to conceptualization, methodology, formal analysis, writing – review & editing, funding acquisition, supervision and project administration.

All authors have read and agreed to the published version of the manuscript.

## Acknowledgments

This research was supported by Fundação para a Ciência e a Tecnologia (FCT, Portugal), project PTDC/SAU-ENB/117013/2010 and Exploratory Project 2022.02763.PTDC. NMS and ASM acknowledge FCT fellowships SFRH/BD/144585/2019 and PD/BD/113698/2015, respectively. MJS acknowledges individual support awarded by FCT Scientific Employment Stimulus (CEECIND/00098/2018). ICM acknowledges consecutive funding from FCT programs “Investigador FCT” (research contract IF/00772/2013) and “Stimulus of Scientific Employment” (research contract CEECIND/01670/2017).

Mass spectrometry data were generated by the UniMS – Mass Spectrometry Unit, iBET/ITQB, Oeiras, Portugal. Authors thank Prof. Manuel Prieto (Instituto Superior Técnico, Universidade de Lisboa, Portugal), Prof. Ana Coutinho (Faculdade de Ciências, Universidade de Lisboa, Portugal) and Prof. Miguel Castanho (Faculdade de Medicina, Universidade de Lisboa, Portugal) for the useful discussions and suggestions.

