## Supplementary data for "Dengue virus capsid protein interaction with nucleic acids"

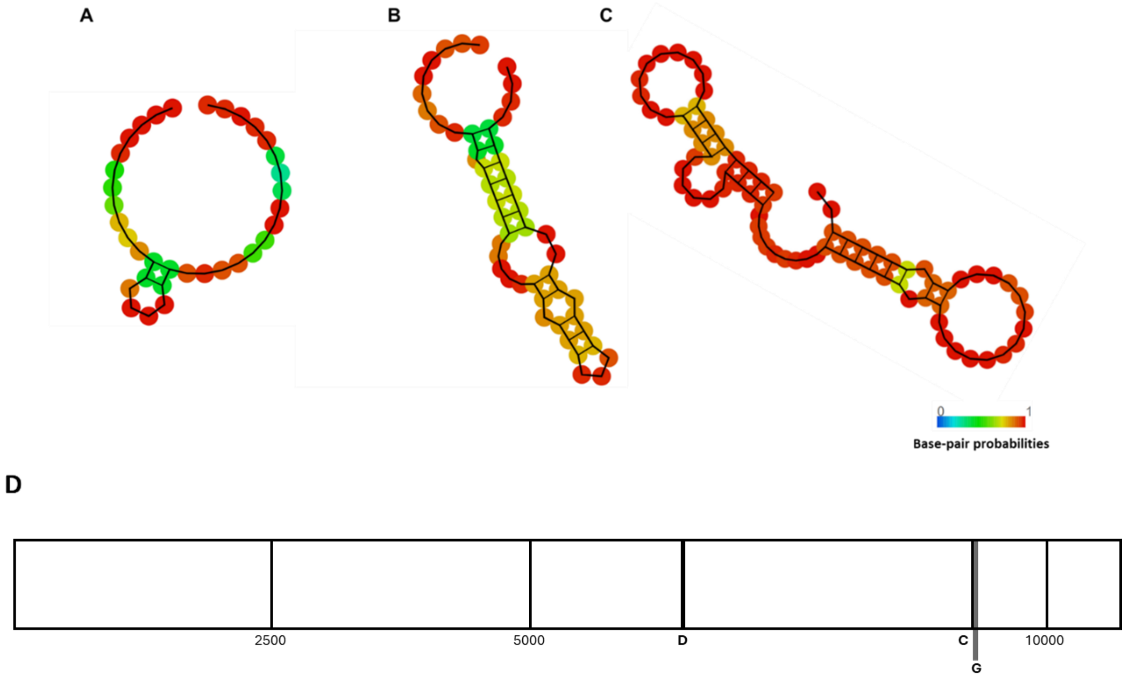


**Figure S1. Characteristics of selected ssRNA sequences.** Secondary structure predictions for sequences with 35 (**A**), 50 (**B**) and 75 (**C**) nucleotides are presented. RNAfold Web Server was used to predict the secondary structures based on the minimum free energy. Structures are colored by base-pair probabilities. Regions of the viral genome analogous to the selected ssDNA sequences (**D**).

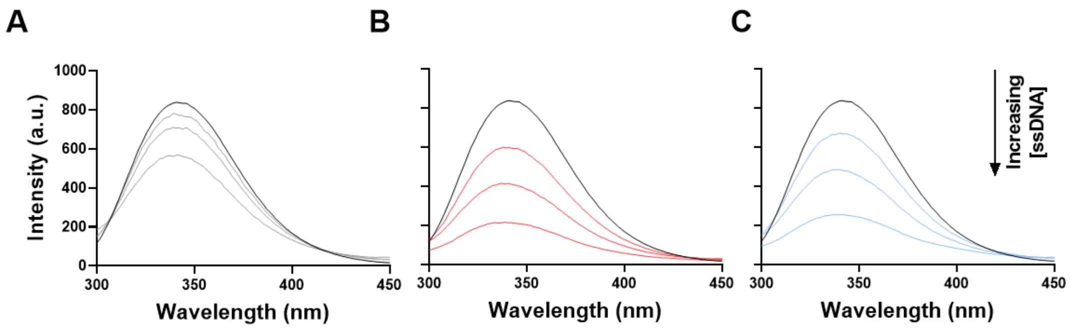


**Figure S2. Fluorescence emission of DENV C in the presence of increasing concentrations of ssDNA with 35 (sequence C) (A), 50 (sequence D) (B) and 75 (sequence G) (C) nucleotides**. The top black line is DENV C emission in the absence of ssDNA. Initial DENV C concentration was kept at 2 μM for all ssDNA sequences. ssDNA concentration varied from 156 nM to 625 nM.

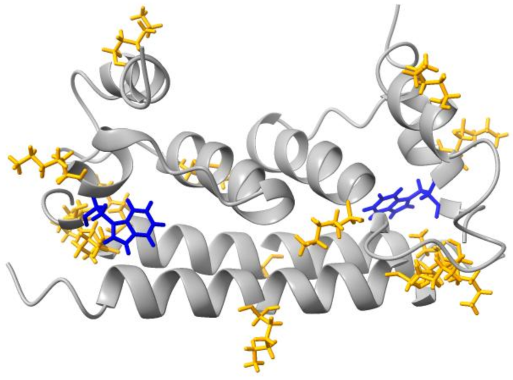

**Figure S3. DENV C structure, highlighting tryptophan (blue) and lysine (orange) amino acid residues.**

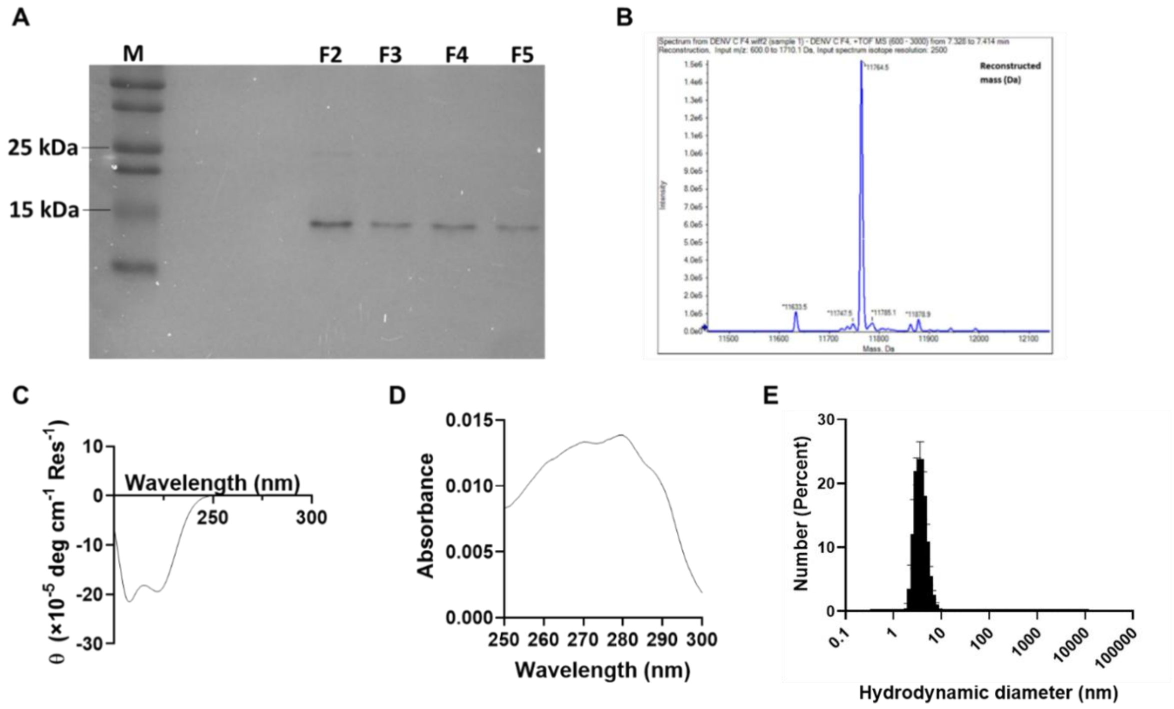


**Figure S4. Protein quality controls.** SDS-PAGE (M, marker; F2-F5, DENV C fractions from size exclusion chromatography) (**A**). Mass spectrum (**B**). CD spectrum**(C)**. UV-Vis absorbance spectrum **(D)**. DLS histogram (particle size number-weighted distribution, in percentage) (**E**).

**Table S1. Characteristics of the single-stranded DNA sequences analogous to regions of the viral genome that are involved in the interaction between DENV C and the viral RNA.**

| Name | Sequence (5’-3’) | Length (mer) | GC content (%) |
|---|---|---|---|
| **C** | AAAGACCAACACCAAGAGGCACAGTAATGGACATT | 35 | 42.9 |
| **D** | GACAACTTAGCAGTCTTGCACACGGCTGAAGCAGGCGGAAGGGCGTACAA | 50 | 56.0 |
| **G** | GTGCGTGTACAAAGACCAACACCAAGAGGCACAGTAATGGACATTATATCGAGAAGAGACCAAA GAGGTAGTGGA | 75 | 45.3 |
| **Scramble C** | AGCAGAACTATACACAGCAGTACAAGGCTACAAGA | 35 | 42.9 |
| **Scramble D** | AGTGCGAACCGCGTCAACACAGGTAGATGGCGGACACATGATTCAAGGCG | 50 | 56.0 |
| **Scramble G** | AGAACGAAGCACGATGAGTATCAGCATTGGTAAGAGCGTAACGAACAACGATCTAGCAGAACAG ATATGGCAGAG | 75 | 45.3 |
| **AT35** | ATTATAAATAATTAATAAAAATTTTATATTAAAAA | 35 | 0.0 |
| **AT75** | ATAATAAAATTATAAAATTAATATAAAAATAATATTTTTATTTAAATAAATTAAAATATAATATTA AAATTATTT | 75 | 0.0 |
| **GC35** | CCGCCGGGTCGGCCGCCGCGGCCGGGGGGGCCCCC | 35 | 100.0 |
| **GC75** | GCGCCCGGGCGGGGGGGGGCCCGGGCCGCGCGGCCCCCGCCCCGGCGCGCGGCGCGCCGCGGGC CGCCCGCCGTC | 75 | 100.0 |

**Table S2. Average maximum emission wavelength at the highest [ssDNA]/[DENV C] ratio.** Results are expressed as mean ± standard deviation from at least 2 independent experiments.

| | Wavelength (nm) |
|---|---|
| **Control** | 341.5 ± 2.1 |
| **C** | 340.0 ± 2.8 |
| **D** | 338.6 ± 1.5 |
| **G** | 337.7 ± 2.1 |
| **Scramble C** | 337.7 ± 1.5 |
| **Scramble D** | 338.6 ± 2.5 |
| **Scramble G** | 339.1 ± 2.1 |
